# Arbuscular cotton-associated mycorrhizal fungi in Yeola region of Maharashtra, India

**DOI:** 10.1101/485326

**Authors:** S W Patale

## Abstract

Mycorrhizae are a mutual symbiotic link between the plant root and a fungus that colonizes the cortical tissue of the roots during active plant growth periods. Both the host plant and the fungus have the potential to benefit. Mycorrhizae are ubiquitous throughout the world in terrestrial ecosystems. The purpose of this study is to evaluate the association of arbuscular mycorrhizal fungi in cotton crops with AM fungal population density in rhizosphere soils, investigate the qualitative composition of AM fungal species and the percentage of root colonization. The results showed that the number of AM fungal propagules collected from different locations in cotton crops ranged from 235 to 1580 spores per 100 g of soil. Due to the widespread nature of AM fungi, they occurred in almost all soil samples, but the number and type of spores and sporocarps varied. In total, 41 AM fungal species belonging to the genera *Glomus*, *Acaulospora* and *Scutellospora* were isolated. *Glomus* was found to be predominantly followed by *Scutellospora* in cotton soils in the rhizosphere. The distribution of spores, density and composition of AM fungi are observed to be influenced by environmental and physicochemical factors. The AM spore number, root colonization percentage and distribution vary depending on the seasonal fluctuations in moisture, temperature, pH and soil mineral nutrient status such as OC, P_2_O_5_, K_2_O, Zn, Cu, Fe, Mn, etc. The obtained data shows that nitrogen-deficient soils had more AM fungal propagules. The soils with a high concentration of phosphorus and potassium had the least AM fungal spores. Depleted zinc, copper and manganese levels have also been positive for more fungal occurrence and distribution. The presence of high iron levels in the soil, however, encourages more AM spores and a percentage of root colonisation.

## INTRODUCTION

Since the Paleozoic era (Taylor 1990), Mycorrhizae has been linked to vascular plants. Arbuscular mycorrhizae (AM), the most prevalent association of plant fungi, comprises about 150 species belonging to the Zygomycotina Glomales (Morton & Bentivenga 1994; Myrold 2000; Perry *et al.*, 1989; Schenk 1981; Simon1996). Arbuscular mycorrhizal (AM) symbiosis is an association between most terrestrial plants and a host plant class of fungi (Glomeromycota) (Schussler *et al.*, 2001). Root colonization with AM fungi has been shown to improve the productivity of many crop plants in dry soils (Al-Karaki and Al-Raddad 1997; Al-Karaki and Clark 1998; Faber *et al.*, 1990 and Sylvia *et al.*, 1993). Ultimately AM fungi improve soil structure by binding soil particles together.

Cotton belongs to the family Malvaceae. It is native to the tropics and warm temperate regions. Commercial species of cotton plant are *G. hirsutum* (90% of world production), *G. barbadense* (8%), *G. arboreum* and *G. herbaceum* (together, 2%).

Nehl *et al.*, (1996, 1998) demonstrated that mycorrhizal colonization, root browning and soil properties associated with cotton growth disorder in Australia and slow mycorrhizal arbuscular colonization in field - grown cotton due to soil environmental conditions. Pattinson *et al.*, (1997) studied the effect of Terrazole and Terraclor fungicides and Fenamiphos nematicide on *Glomus* mosseae root colonization and the growth of cotton seedlings. Zak *et al.*, (1998) studied Arbuscular-mycorrhizal colonization dynamics of cotton (*Gossypium hirsutum* L.) growing under several production systems on the Southern High Plains of Texas. Mc Gee, *et al.*, (1999) have reviewed the relationship between density of *Glomus mosseae* propagules and the initiation and spread of arbuscular mycorrhizas in cotton roots. Feng *et al.*, (2002) have shown the uptake of nitrogen from indigenous soil pool by cotton plant inoculated with arbuscular mycorrhizal fungi. Hulugalle *et al.*, (2004) studied soil properties, and cotton growth, yield and fiber quality in three cotton-based cropping systems. Patale and Shinde (2012b) studied effect of water stress on growth performance of Bt% cotton inoculated with AM Fungi. Patale and Shinde (2012a) described growth performance of Bt% cotton inoculated with AM Fungi on salinity stress. Patale and Shinde (2012c) studied influence of *Glomus* species and soil phosphorous on *Verticillium* wilt in Bt% cotton.

Cotton is cultivated in Maharashtra as a chief commercial crop, survey of literature do not show any report on association of AM Fungi with cotton in this area. The study of this association will definitely be useful to all cotton growers to increase yield and to improve fertility of soil. The use of AM fungi will also reduce soil pollution due to application of Chemical fertilizers and fungicides in cotton fields as AM fungi are ecofriendly.

## MATERIALS AND METHODS

In the present study, cotton crop commonly grown, in and around Yeola viz., Saigaon (S1), Patoda (S2), Golhewadi (S3), Kotamgaon (S4), Bharam (S5), Kusur (S6), Gawandgaon (S7) and Mukhed (S8) were surveyed for arbuscular mycorrhizal (AM) fungal association.

Fresh samples of soil were taken to the laboratory. Fine roots were fixed in the formalin acetic acid alcohol solution (90:5:5) after washing thoroughly to determine the root infection. Soil samples were dried air in the shade for additional spores at laboratory temperature. Roots were autoclaved in KOH solution for 15 to 20 minutes (10 per cent), cleaned in distilled water and neutralized with HCl (2 per cent) and stained in lacto phenol in trypan blue (0.05 per cent). Phillips and Hayman's (1970) method measured the percentage of the root infection.

100 g of air - dried soil mixture with 1000 ml of tap water has been placed in a beaker. The mixture of the root soil has been strongly mixed with glass rod for 30 seconds. The remaining soil-root-hyphae-spore suspension was slowly poured through 240, 170, 150,100 and 72 μm sieves after the soil particles and organic debris were settled. The extracts were washed off the sieves to what man filter paper. Spores, aggregates and sporocarps were picked by needle using trinocular research microscopes (Gerdemann and Nicolson, 1963). To each PVLG drop, 5 - 10 spores have been added. The mounting system was allowed to set for 3 – 5 minutes before a cover slip was added. The identification of isolated spores was carried out by Schenck and Perez (1990) using the key proposed.

The cotton rhizosphere soil was analyzed for the physical and chemical characteristics of soil samples such as pH, EC, OC, P, K, Zn, Cu, Fe and Mn performed using Jackson's (1973) procedure.

## RESULTS AND DISCUSSION

### Ecology of AM fungi

The presence of AM fungi was screened for cotton plants belonging to the Malvaceae family. Data on fungal propagules isolated from these plants ' soil samples collected from different locations are given in Table 1. Soil samples collected from eight different locations were found to be associated with AM fungus. All samples collected from the ground showed a percentage of root colonization, regardless of location (Table 1). The data indicates the number of AM fungal propagules accumulated in different crops from different locations ranging from 235 to 1580 spores per 100 g soil. Of all the 8 soil samples surveyed, Mukhed soils contain more AM propagules of 1580 per 100 g of soil; while Patoda soil samples showed less than 235 spores per 100 g of soil. In all, 41 AM fungal species representing three genera, namely *Acaulospora*, *Glomus* and *Scutellospora*, have been isolated (Table 2).*Glomus* representing twenty - two species, seven species of *Acaulospora* and twelve species of *Scutellospora.* Out of these twenty AM fungal species, the present soils are dominated by *Glomus* species followed by *Scutellospora*.

### Physico-chemical factors of the Soil

Table 1 presents data on the physico-chemical characteristics of the rhizosphere soil samples of cotton collected from different locations in relation to the number of propagules. All the soils investigated in this study were of sandy loam type and were fertilized organically and chemically. The soils had a pH range of 8 to 8.95. More propagules were shown in the slightly alkaline pH soils.

Data from the soils of the rhizosphere showed that organic carbon-deficient soils had more AM fungal propagules. Soils with a low organic carbon content of 0.22% had 1580 AM fungal propagules per 100 g. While high organic carbon soil samples (1.23 percent) had a lower spore density of 240 per 100 g. These soil samples were studied for their phosphorus content and it was observed that soils containing 3 to 4 kg / ac phosphorus had a maximum number of AM fungal spores, while soils with a lower number of AM fungal spores had high phosphorus content, i.e. 12 to 16 kg per acre.

High potassium levels contain the smallest amount of AM spores compared to soils with low potassium levels. It was therefore evident that low potassium levels favored more AM fungal spore association. Similarly, soils with minimum levels of copper, zinc, iron and manganese were favorable for the occurrence and distribution of more AM funguses. However, high iron levels are favorable for more AM spore occurrence.

### AM fungal root colonization

The percentage of AM cotton fungal root colonization from all selected sites is shown in table 1. It is clear from the data that the percentage of root colonization in the cotton samples collected at different locations varied. The percentage of root colonization in the Mukhed samples was found to be maximum (95%) and minimum (63.64%) in Patoda samples. Saigaon root samples were found to be heavily colonized with vesicles.

In this study, the population dynamics of AM fungi were determined by the collection of the remaining spores in and around Yeola from different soils of brinjal, tomato and cotton. Because of the widespread nature of AM fungi, these occurred in nearly all soil samples, but with a variation in the number and type of spores and sporocarps regardless of the location.

A total of 41 AM fungal species have been isolated from eight different soils of *Glomus, Acaulospora* and *Scutellospora* genera. Among the soil samples collected from Saigaon, there were more AM propagules followed by Mukhed soil samples, which could be due to low nutrient status and average humidity levels.

*Glomus* was predominantly among the 41 AM fungal species isolated from the rhizospheric soils of cotton crops, followed by *Scutellospora*. The results support the previous studies very strongly (Patale and Shinde 2010a, b; Patale 2016). Earlier reports also revealed the predominance of the above-mentioned AM fungal genera in plant cultivar rhizosphere soils (Gerdemann and Trappe, 1974; Hall and Abbott, 1984).

### Effect of soil's physico-chemical factors

It has been observed that the distribution of spores, density and composition of AM fungi are influenced by environmental and physicochemical factors. The number of AM spores, the percentage of root colonization and the distribution are affected by the seasonal fluctuations in humidity, temperature, pH and soil nutrient status such as N, P, K, Zn, Fe, etc. Earlier studies in chili, sorghum, mungbean, tomato, brinjal and soybean by Bagyaraj and Sreeramulu (1982), Reddy *et al.*, (2006), Reddy *et al.*, (2007), Patale and Shinde (2010a, b) and Patale (2016) also showed similar trends.

In the present study, the population of mycorrhizal spores in rhizosphere soil and the percentage of mycorrhizal infection in plant roots fluctuated with changes in soil physico-chemical factors. The results coincide with the earlier results of Reddy *et al.*, (2007) in sorghum crops.

Light textured sandy loam soil with neutral to slightly alkaline pH, low humidity favored an extensive association of mycorrhizal roots (Sreeramulu and Bhagyaraj, 1986). The pH of the soil in our study ranged from 8 to 8.95, which was slightly alkaline and had more, AM fungal propagules. In this study, soils with low organic carbon levels such as 0.17 percent of the soil contained 1278 AM fungal propagules per 100 gm of soil and 92 percent root colonization, but soil samples with a nitrogen content of 1.23 percent contained only 240 spores per 100 g of soil and 85.71 percent root colonization. It therefore clearly indicates that high nitrogen levels reduce the number of AM propagules and the percentage of root infection that is in line with the research published by Azcon-Aquilar and Barea (1982), Patale and Shinde (2010a, b) and Patale (2016).

The deficiency of phosphorus in semi - arid soils is the rule of the association of AM plants (Williams *et al.*, 1974). Mosse (1981) observed that high levels of free soil phosphorus decreased mycorrhizal development, while Ojala *et al.*, (1983) recorded a decrease in plant mycorrhizal dependency with an increase in phosphorus available in the soil. Our study also showed the same trend that samples with a higher phosphorous content (16 kg / ac) had less AM propagules than the soil with a lower phosphorus content (3 to 4 kg / ac) with a maximum number of AM spores. AM fungi, particularly phosphorus (P), have been found to improve plant mineral nutrition. Smith and Read (1997) obtained the same results.

According to our experimental results of the soil samples, high potassium levels in the soil had less AM spores than those with low potassium levels. Our findings coincide with Barea *et al.*, (2002), and Suresh *et al.*, (2000)’s observations that low potassium levels favored more AM fungal associations than high potassium levels in the soil. Similar trends have been observed that low concentrations of Zn, copper and manganese and high iron levels favor AM fungal propaganda. These comments are consistent with the previous reports (Barea et al., 2002; Brundrett, 2002; Garg and Chandel, 2010).

In the present study, the genus *Glomus* has been represented by more species than the other genera, which indicates its predominance. This study gives a clear understanding of the AM fungi ecology.

Because of the widespread nature of AM fungi, these occurred in nearly all soil samples, but with a variation in the number and type of spores and sporocarps regardless of the location. The *Glomus* genus was represented by more species than the other genera, which showed its predominance. This study gives a clear understanding of the AM fungi ecology.

It was concluded that AM fungi were adapted to crop plants and the environment that manage indigenous fungal populations in agricultural practices so that the population of efficient indigenous fungi is increased.

**Table. 1.**
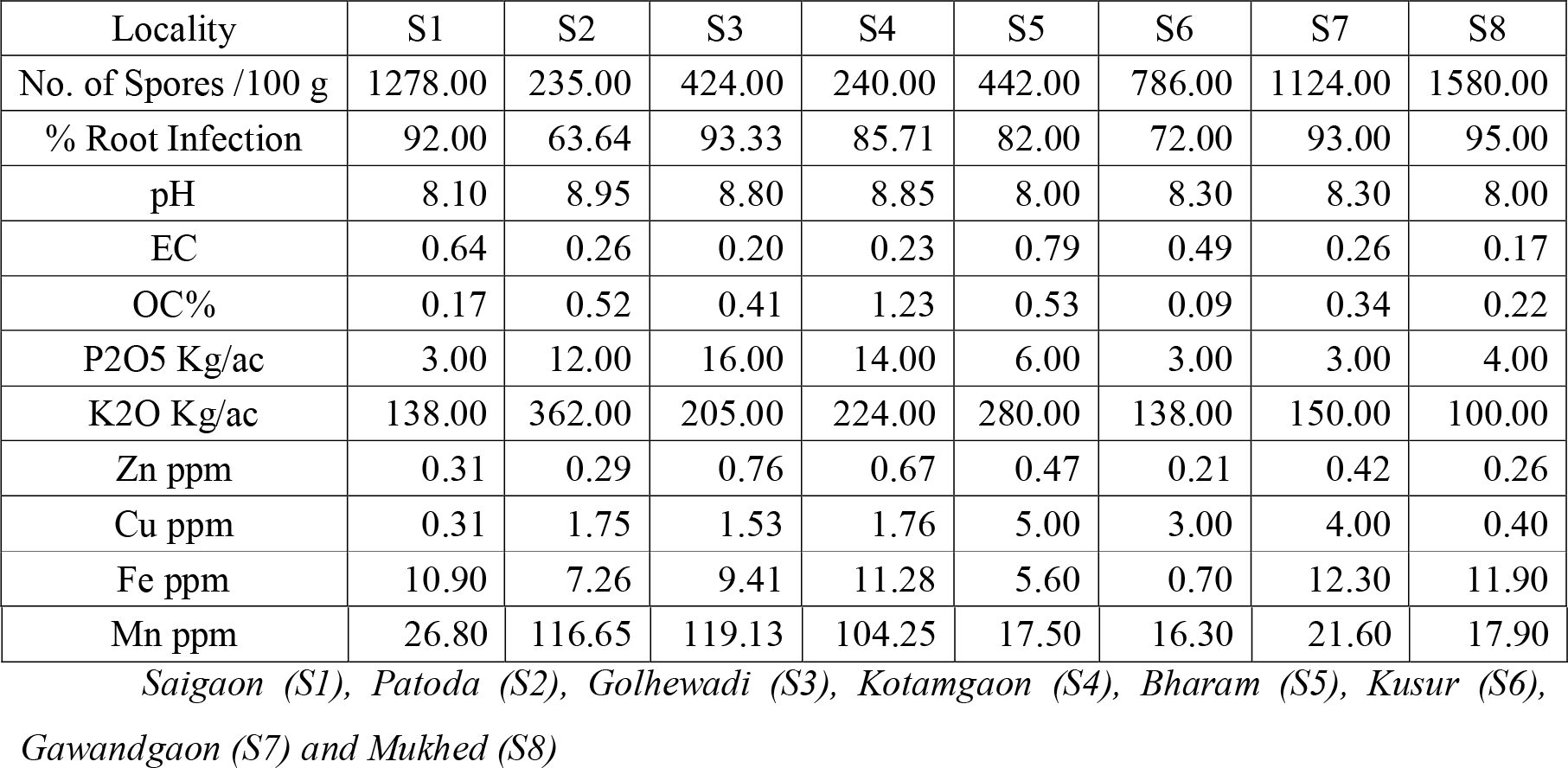
Number of AM fungal propagules in relation to physico-chemical factors in the rhizosphere soil samples of Cotton

| Locality | S1 | S2 | S3 | S4 | S5 | S6 | S7 | S8 |
| --- | --- | --- | --- | --- | --- | --- | --- | --- |
| No. of Spores /100 g | 1278.00 | 235.00 | 424.00 | 240.00 | 442.00 | 786.00 | 1124.00 | 1580.00 |
| % Root Infection | 92.00 | 63.64 | 93.33 | 85.71 | 82.00 | 72.00 | 93.00 | 95.00 |
| pH | 8.10 | 8.95 | 8.80 | 8.85 | 8.00 | 8.30 | 8.30 | 8.00 |
| EC | 0.64 | 0.26 | 0.20 | 0.23 | 0.79 | 0.49 | 0.26 | 0.17 |
| OC% | 0.17 | 0.52 | 0.41 | 1.23 | 0.53 | 0.09 | 0.34 | 0.22 |
| P2O5 Kg/ac | 3.00 | 12.00 | 16.00 | 14.00 | 6.00 | 3.00 | 3.00 | 4.00 |
| K2O Kg/ac | 138.00 | 362.00 | 205.00 | 224.00 | 280.00 | 138.00 | 150.00 | 100.00 |
| Zn ppm | 0.31 | 0.29 | 0.76 | 0.67 | 0.47 | 0.21 | 0.42 | 0.26 |
| Cu ppm | 0.31 | 1.75 | 1.53 | 1.76 | 5.00 | 3.00 | 4.00 | 0.40 |
| Fe ppm | 10.90 | 7.26 | 9.41 | 11.28 | 5.60 | 0.70 | 12.30 | 11.90 |
| Mn ppm | 26.80 | 116.65 | 119.13 | 104.25 | 17.50 | 16.30 | 21.60 | 17.90 |
| Saigaon (S1), Patoda (S2), Golhewadi (S3), Kotamgaon (S4), Bharam (S5), Kusur (S6),<br>Gawandgaon (S7) and Mukhed (S8) |  |  |  |  |  |  |  |  |

**Table. 2.**
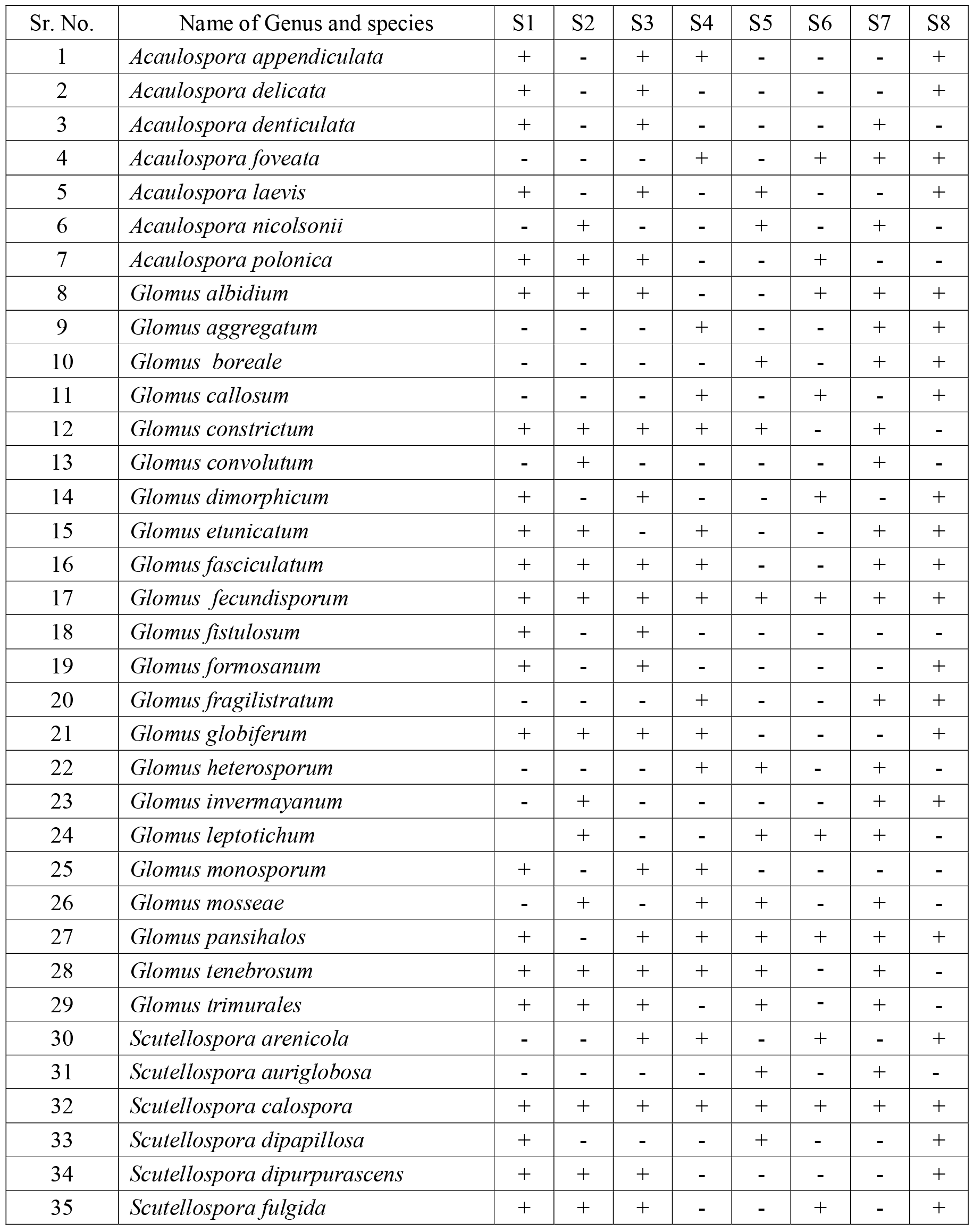
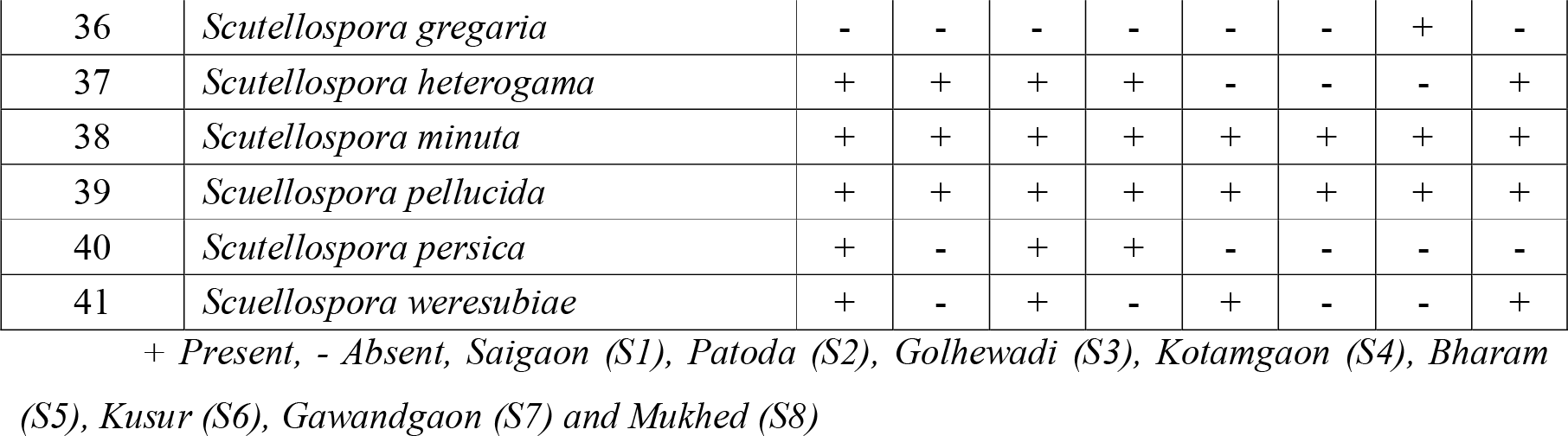
Arbuscular mycorrhizal fungal spore distribution in the rhizosphere soils of Cotton collected from different locations

| Sr. No. | Name of Genus and species | S1 | S2 | S3 | S4 | S5 | S6 | S7 | S8 |
| --- | --- | --- | --- | --- | --- | --- | --- | --- | --- |
| 1 | *Acaulospora appendiculata* | + | - | + | + | - | - | - | + |
| 2 | *Acaulospora delicata* | + | - | + | - | - | - | - | + |
| 3 | *Acaulospora denticulata* | + | - | + | - | - | - | + | - |
| 4 | *Acaulospora foveata* | - | - | - | + | - | + | + | + |
| 5 | *Acaulospora laevis* | + | - | + | - | + | - | - | + |
| 6 | *Acaulospora nicolsonii* | - | + | - | - | + | - | + | - |
| 7 | *Acaulospora polonica* | + | + | + | - | - | + | - | - |
| 8 | *Glomus albidum* | + | + | + | - | - | + | + | + |
| 9 | *Glomus aggregatum* | - | - | - | + | - | - | + | + |
| 10 | *Glomus boreale* | - | - | - | - | + | - | + | + |
| 11 | *Glomus callosum* | - | - | - | + | - | + | - | + |
| 12 | *Glomus constrictum* | + | + | + | + | + | - | + | - |
| 13 | *Glomus convolutum* | - | + | - | - | - | - | + | - |
| 14 | *Glomus dimorphicum* | + | - | + | - | - | + | - | + |
| 15 | *Glomus etunicatum* | + | + | - | + | - | - | + | + |
| 16 | *Glomus fasciculatum* | + | + | + | + | - | - | + | + |
| 17 | *Glomus fecundisporum* | + | + | + | + | + | + | + | + |
| 18 | *Glomus fistulosum* | + | - | + | - | - | - | - | - |
| 19 | *Glomus formosanum* | + | - | + | - | - | - | - | + |
| 20 | *Glomus fragilistratum* | - | - | - | + | - | - | + | + |
| 21 | *Glomus globiferum* | + | + | + | + | - | - | - | + |
| 22 | *Glomus heterosporum* | - | - | - | + | + | - | + | - |
| 23 | *Glomus inermayanum* | - | + | - | - | - | - | + | + |
| 24 | *Glomus leptotichum* |  | + | - | - | + | + | + | - |
| 25 | *Glomus monosporum* | + | - | + | + | - | - | - | - |
| 26 | *Glomus mosseae* | - | + | - | + | + | - | + | - |
| 27 | *Glomus pansihalos* | + | - | + | + | + | + | + | + |
| 28 | *Glomus tenebrosus* | + | + | + | + | + | - | + | - |
| 29 | *Glomus trimurales* | + | + | + | - | + | - | + | - |
| 30 | *Scutellospora arenicola* | - | - | + | + | - | + | - | + |
| 31 | *Scutellospora auriglobosa* | - | - | - | - | + | - | + | - |
| 32 | *Scutellospora calospora* | + | + | + | + | + | + | + | + |
| 33 | *Scutellospora dipapillosa* | + | - | - | - | + | - | - | + |
| 34 | *Scutellospora dipurpurascens* | + | + | + | - | - | - | - | + |
| 35 | *Scutellospora fulgida* | + | + | + | - | - | + | - | + |
| 36 | *Scutellospora gregaria* | - | - | - | - | - | - | + | - |
| 37 | *Scutellospora heterogama* | + | + | + | + | - | - | - | + |
| 38 | *Scutellospora minuta* | + | + | + | + | + | + | + | + |
| 39 | *Scutellospora pellucida* | + | + | + | + | + | + | + | + |
| 40 | *Scutellospora persica* | + | - | + | + | - | - | - | - |
| 41 | *Scutellospora weresubiae* | + | - | + | - | + | - | - | + |

+ Present, - Absent, Saigaon (S1), Patoda (S2), Golhewadi (S3), Kotamgaon (S4), Bharam (S5), Kusun (S6), Gawandgaon (S7) and Mukhed (S8)

